# Foxp1 Coordinates Early Temporal Identity and Neurogenic Progression in Retinal Progenitors

**DOI:** 10.64898/2026.10.23.760569

**Authors:** Fei Chang, Jianmin Zhang, Jacqueline M. Roberts, Joon Schwakopf, Michel Cayouette, Joel B. Miesfeld, Pierre Mattar, Monica L. Vetter

## Abstract

Neural diversity depends on both the temporal competence of progenitors to generate particular cell types and their progression into a neurogenic state. How these distinct features of progenitor development are mechanistically coordinated remains poorly understood. Here, we identify Foxp1 as a dual function transcriptional regulator that couples early progenitor identity to neurogenic progression in the developing retina. By defining its direct transcriptional targets, we show that Foxp1 coordinates complementary regulatory programs: it sustains early competence by activating early-stage retinal progenitor genes and repressing late temporal factors, while promoting neurogenic progression through activation of the proneural factors *Atoh7* and *Neurog2*. We establish a cis-regulatory mechanism linking these functions: Foxp1 activates *Atoh7* through its remote shadow enhancer, and deletion of this enhancer prevents Foxp1-dependent induction of *Atoh7* and retinal ganglion cell (RGC) production. We further reveal that Neurog2 partially compensates for loss of Atoh7 function. This coordinated early progenitor program is temporally restricted by progressive silencing of *Foxp1*, involving sequential deposition of H2AK119ub1 followed by H3K27me3 at a distal conserved silencer essential for its repression. Together, these findings establish a mechanistic link between temporal competence and neurogenic progression, revealing how a shared transcriptional regulator coordinates progenitor identity with neuronal production through distinct cis-regulatory elements.

**Significance statement:** Temporal competence and neurogenic progression are distinct features of neural progenitor development, yet how they are mechanistically coordinated remains poorly understood. We identify Foxp1 as a transcriptional regulator that couples early progenitor identity to neurogenic progression in the developing retina. Foxp1 activates early competence genes, represses late temporal factors, and directly activates proneural regulators. Functional dissection of cis-regulatory elements establishes an enhancer requirement for Foxp1-dependent neuronal production and a silencer requirement for repression of Foxp1 itself. These findings reveal how coordinated transcriptional activation and repression connect progenitor competence to neuronal differentiation, providing a mechanistic framework for understanding the temporal control of neural cell diversity.

## Introduction

Central nervous system (CNS) function relies on the generation of diverse neuronal types at the appropriate developmental times and in precise proportions. Multipotent neural progenitor cells progress through temporal competence states that shape the neuronal types they can produce. Intrinsic programs and temporal transcription factors regulate these states across different CNS regions (1, 2). Within each temporal state, subsets of progenitors activate proneural basic helix-loop-helix (bHLH) factors and enter a neurogenic program that initiates neuronal differentiation (3, 4). Temporal competence and neurogenic progression thus represent distinct, but coordinated, features of progenitor development: one defines the range of available neuronal fates, whereas the other governs progression toward their production. Identification of temporal regulators and proneural factors have advanced our understanding of each process (2, 4), but how their transcriptional programs are mechanistically connected remains poorly understood. Resolving this connection is essential to explain how progenitor identity is translated into the production of developmentally appropriate neurons.

The developing retina provides a tractable system for resolving this relationship. Retinal progenitor cells (RPCs) pass through distinct competence states to sequentially generate early- and late-born retinal cell types (5). Single cell sequencing studies distinguish rapidly dividing primary RPCs from neurogenic RPCs, which are enriched for genes encoding proneural bHLH factors and give rise to retinal neurons (6-11). Both populations undergo temporal changes in gene expression as retinal cell production shifts from early- to late-born types. Regulators of primary RPC temporal competence include the early factors *Foxp1*, *Ikzf1*, and *Pou2f1/2*, and the mid/late factors *Casz1* and *Nfia/b/x* (5, 10, 12-14). Neurogenic RPCs also express temporally patterned proneural genes, with *Atoh7* and *Neurog2* enriched in early subsets and *Ascl1* enriched in later subsets (10). Thus, temporal identity and entry into a neurogenic state represent distinguishable aspects of RPC development, providing an opportunity to define the regulatory mechanisms that coordinate these transitions.

Foxp1 is a candidate molecular link between these programs. We previously established that Foxp1 regulates early RPC competence and early-born retinal neuron production, and that Jarid2-dependent repression of *Foxp1*, associated with H3K27me3 deposition, enables the transition to late RPC competence (5). In the developing cerebral cortex, Foxp1 maintains radial glial identity and regulates the competence window for producing early-born, deep-layer neurons (15). Studies in human cortical organoids and mouse cortex implicate Foxp1 in regulating the balance between progenitor proliferation and neuronal differentiation (16, 17). These findings support roles for Foxp1 in progenitor identity and neurogenesis across CNS regions and species, raising the possibility that it coordinates temporal competence with neurogenic progression to govern the production of developmentally appropriate neurons.

Here, we show that Foxp1 acts as a dual function transcriptional regulator that couples early progenitor identity to neurogenic progression. By defining its direct targets, we show that Foxp1 activates early-stage primary RPC genes and represses late temporal factors, while directly activating the proneural genes *Atoh7* and *Neurog2* to promote entry into a neurogenic state. We establish a mechanistic basis for Foxp1-dependent neuronal production, showing that activation of *Atoh7* requires its remote shadow enhancer (SE) (18), while deletion of this enhancer prevents Foxp1-dependent induction of *Atoh7* and retinal ganglion cell (RGC) production. We further reveal that Neurog2 partially compensates for loss of Atoh7 function, promoting incomplete RGC generation. Finally, we find that restriction of this early program through progressive silencing of *Foxp1* involves sequential deposition of H2AK119ub1 followed by H3K27me3 and requires a conserved distal silencer element. Together, these findings establish a mechanistic connection between progenitor temporal identity and neurogenic progression and reveal how cis-regulatory control temporally coordinates this program.

## Results

### *Foxp1* directly activates early RPC genes and represses late RPC genes

To investigate how *Foxp1* regulates gene expression to maintain early temporal patterning of RPCs and promote the generation of early-born retinal neurons, we mapped its genomic binding sites. CUT&RUN for Foxp1 was performed with wild type retina at E14.5, when Foxp1 has high endogenous expression. To add robustness, we also performed this experiment in *Six3-Cre; Foxp1*^tg/+^ (*Foxp1* cTG) retina at P0, where Foxp1 transgene expression is sustained in RPCs resulting in extended early RPC competence and increased production of early-born cell types (5). By overlapping Foxp1 peaks identified in both endogenous and transgenic conditions, we identified 3,753 high-confidence Foxp1 consensus peaks (Figure 1A and 1B). De novo motif analysis using HOMER revealed a significant enrichment of the Forkhead family motif within Foxp1 binding sites (Figure 1C).

**Figure 1:**
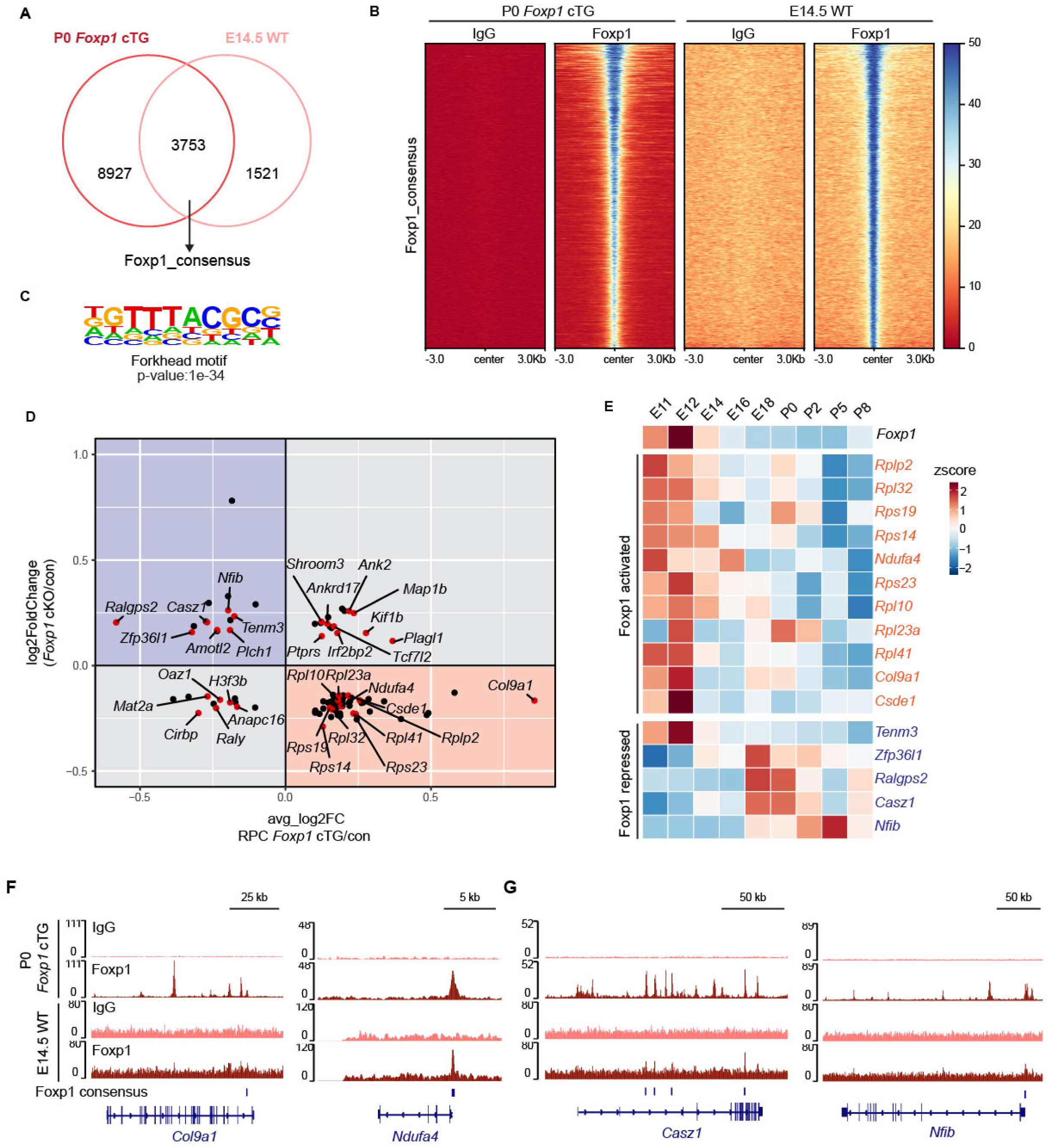
Foxp1 maintains early RPC state by directly activating early RPC genes and repressing late RPC genes. (A) Venn diagram showing Foxp1 peaks identified by CUT&RUN in P0 *Six3*-Cre; *Foxp1*^tg/+^ (*Foxp1* cTG) and E14.5 WT retina. Foxp1 consensus peaks represent peaks shared between both conditions. (B) Heatmap showing the signal intensities of IgG and Foxp1 centered at Foxp1 consensus peaks (± 3 kilobases (kb)) in P0 *Foxp1* cTG and E14.5 WT retina. (C) Forkhead motif identified by HOMER motif analysis using all Foxp1 consensus peaks; the enrichment *P* value is indicated. (D) Intersection of differentially expressed genes in E16.5 *Six3*-Cre; *Foxp1*^fl/fl^ (*Foxp1* cKO) retina and in P0 *Foxp1* cTG primary RPCs. Genes in the bottom right (orange) quadrant are activated by *Foxp1* and genes in the upper left (blue) quadrant are repressed by *Foxp1*. Genes near Foxp1 consensus peaks are labeled and colored red. (E) Heatmap showing dynamic expression of Foxp1-activated (orange) and Foxp1-repressed (blue) genes in RPCs throughout retinal development (10). (F, G) Foxp1 and IgG CUT&RUN genome tracks at representative (F) early RPC genes (*Col9a1* and *Ndufa4)* and (G) late RPC genes (*Casz1* and *Nfib).* Blue bars below the tracks indicate identified Foxp1 consensus peaks.

We previously showed that in E16.5 *Foxp1* conditional knockout (cKO) retina there was reduction of early RPC genes and increased expression of late RPC genes, with the opposite evident in RPC clusters from scRNA-seq analysis of P0 *Foxp1* cTG retina (5). To identify genes directly regulated by *Foxp1* in early-stage RPCs, we intersected differentially expressed genes (DEGs) from these two transcriptional datasets with genes associated with Foxp1 consensus peaks (Figure 1D). This analysis identified a group of genes directly repressed by *Foxp1*: bound by Foxp1, upregulated in *Foxp1* cKO retina, and downregulated in *Foxp1* cTG RPCs (Figure 1D). Notably, most *Foxp1*-repressed genes are selectively expressed in RPCs at later developmental stages, when *Foxp1* expression is low, including established late temporal identity genes *Casz1* and *Nfib* (Figure 1E and 1G). Conversely, we identified *Foxp1*-activated genes that were bound by *Foxp1.* In accordance with the transient expression of Foxp1 in early RPCs, many of these genes are highly expressed in RPCs during early embryonic stages, including *Col9a1* and *Ndufa4* (Figure 1D, 1E, and 1F). Together, these findings demonstrate that Foxp1 plays a dual role in maintaining the early RPC state by directly activating early RPC genes while simultaneously repressing genes that are important for late temporal identity.

### Foxp1 positively regulates Atoh7 expression

Before differentiating into retinal neurons, RPCs go through a transient neurogenic state marked by the expression of proneural bHLH factors (10). To understand whether *Foxp1* plays a role in regulating this transition, we examined the neurogenic RPC cluster in our P0 *Foxp1* cTG scRNA-seq dataset (5). We found that the neurogenic specific bHLH factor *Atoh7* was significantly upregulated in *Foxp1* cTG neurogenic RPCs and downregulated in E16.5 *Foxp1* cKO retina, suggesting a positive regulatory relationship between *Foxp1* and *Atoh7* (Figure 2A and 2B).

**Figure 2:**
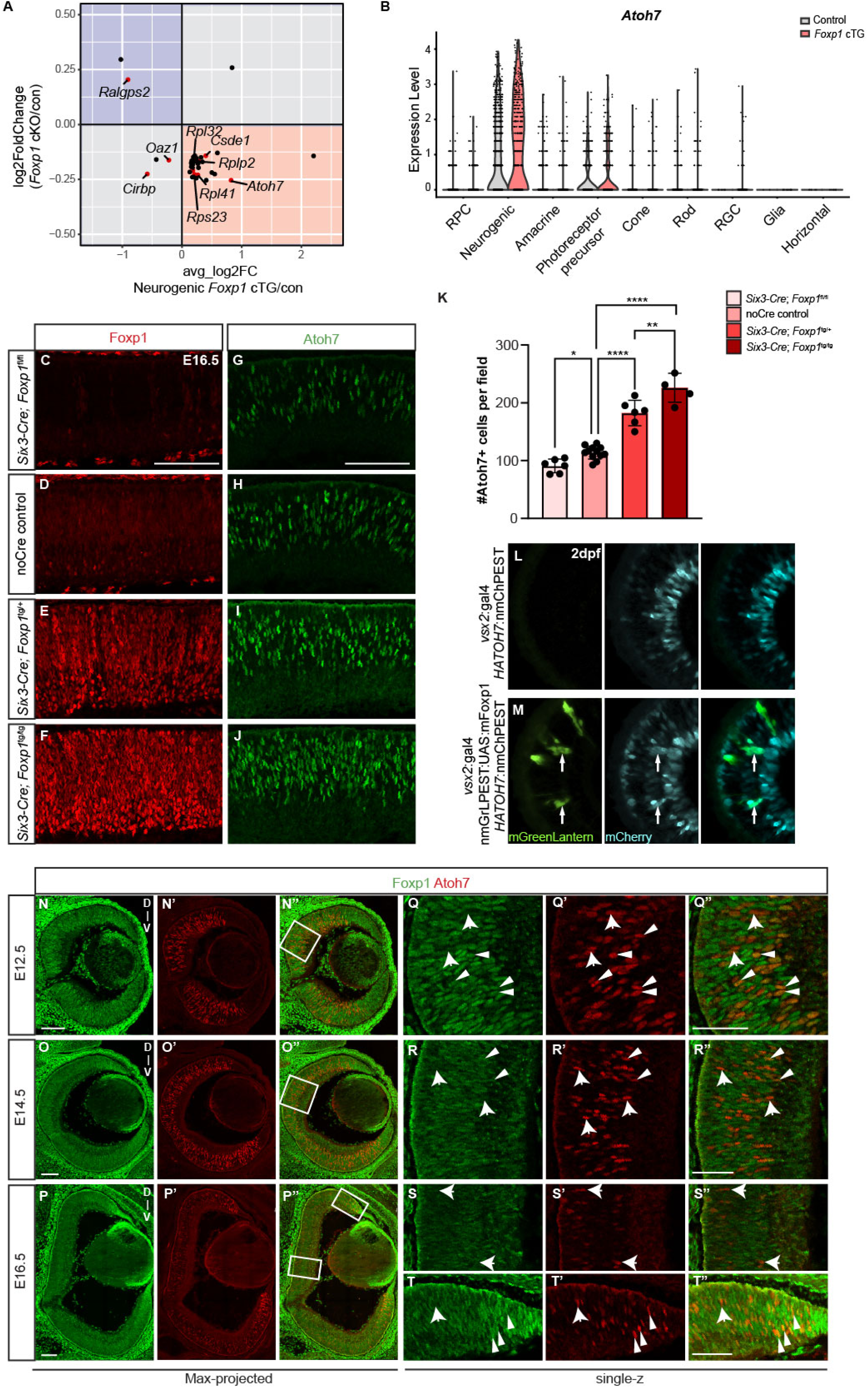
Foxp1 activates Atoh7 through binding to the shadow enhancer. (A) Intersection of differentially expressed genes in E16.5 *Foxp1* cKO retina and in P0 *Foxp1* cTG neurogenic RPCs. Genes in the bottom right (orange) quadrant are activated by *Foxp1*; genes near *Foxp1* consensus peaks are labeled and colored red. (B) Violin plot of *Atoh7* expression in major clusters from P0 control and *Foxp1* cTG scRNA-seq data. (C-J) Representative images showing Foxp1 and Atoh7 expression in the central retina at E16.5 in (C, G*) Six3-Cre*, *Foxp1*^fl/fl^, (D, H) no-Cre control, (E, I) *Six3-Cre*; *Foxp1*^tg/+^ and (F, J) *Six3-Cre*; *Foxp1*^tg/tg^. Scale bar: 100 μm. (K) Quantification of Atoh7+ cells per field within central dorsal retina in different genotypes (*Six3-Cre*, *Foxp1*^fl/fl^: n = 6 retina from 3 animals; no-*Cre* Control: n =12 retina from 6 animals; *Six3-Cre*; *Foxp1*^tg/+^: n = 6 retina from 3 animals; *Six3-Cre*; *Foxp1*^tg/tg^: n = 4 retina from 4 animals). Graphs represent mean ± SEM. *p=0.0361, **p=0.0014, ****p<0.0001 by one-way ANOVA with Sidak’s multiple comparisons. (L, M) Representative images of 2 days post fertilization (dpf) retinas expressing a human ATOH7 BAC transgenic mCherry reporter without (L) or with (M) mouse Foxp1 expression in the *vsx2*-expressing RPC lineage, as indicated by co-expression of mGreenLantern. Arrows indicate cells co-expressing Foxp1-mGreenLantern and the ATOH7-mCherry transgenic reporter. (N-T) Representative confocal images showing Foxp1 and Atoh7 expression in (N-N’’, Q-Q’’) E12.5, (O-O’’, R-R’’) E14.5 and (P-P’’, S-S’’, T-T’’) E16.5 WT mouse retina. (N-P) Max projected images of the whole retinal cross-section; (Q-T) single-z stack (0.8 μm thickness) images of central dorsal (Q-S) and peripheral (T) regions at higher magnification. Arrowheads indicate Foxp1 and Atoh7 double-positive cells; arrows indicate Atoh7 only cells. Scale bar: 100 μm in N-P; 50 μm in Q-T. D-V (Dorsal-Ventral) represents the orientation of the section.

To validate this observation from the sequencing data, we performed Atoh7 immunostaining on E16.5 retinal sections. We generated a series of mouse lines with varying *Foxp1* gene dosage in RPCs: *Foxp1* cKO (*Six3*-Cre; *Foxp1^fl^*^/fl^), endogenous low *Foxp1* level (no-Cre control), one *Foxp1* transgene copy (*Six3*-Cre; *Foxp1*^tg/+^), and two transgene copies (*Six3*-Cre; *Foxp1*^tg/tg^). As *Foxp1* gene dosage increased, Foxp1 protein levels correspondingly rose (Figure 2C-2F). Compared to no-Cre control retina, we found Atoh7+ cell numbers were reduced in *Foxp1* cKO retina, elevated in *Six3*-Cre, *Foxp1*^tg/+^ retina and further increased in *Six3*-Cre, *Foxp1*^tg/tg^ retina (Figure 2G-2K), indicating that Atoh7 expression is sensitive to *Foxp1* levels.

To test if *Atoh7* can be directly activated by *Foxp1* expression, we used a human ATOH7 transgenic zebrafish reporter line (*HATOH7*:nmChPEST), which contains the full ATOH7 regulatory sequence, including the proximal primary enhancer (PE) and remote SE, driving expression of destabilized nuclear mCherry during zebrafish retinal neurogenesis. We expressed mGreenLantern-labeled mouse Foxp1 in the *vsx2*-expressing RPC lineage using *vsx2*:Gal4;nmGrLPEST:*UAS*:mFoxp1. In the mFoxp1 expressing line ectopic mCherry was observed in mGreenLantern+ cells, demonstrating the ability of Foxp1 to induce expression via *Atoh7* regulatory elements and highlighting the functional conservation of Foxp1 between species (Figure 2L-2M).

### Foxp1 and Atoh7 are transiently co-expressed during mouse retinal development

If Foxp1 regulates Atoh7 expression to drive neurogenic progression, we predict overlap in their expression during mouse retinal development, so we examined their spatiotemporal expression by performing immunostaining. Consistent with previous observations, we found that Foxp1 and Atoh7 were dynamically expressed in the developing mouse retina from E12.5 to E16.5 (5, 19). Foxp1 was broadly expressed in progenitors throughout the retina at E12.5 whereas Atoh7 expression was mostly restricted to the central region (Figure 2N-2N’’). At this stage, most Atoh7+ cells co-expressed Foxp1, suggesting that a subset of Foxp1+ RPCs transition into a neurogenic state (Figure 2Q-2Q’’). By E14.5, Foxp1 expression in the central region was slightly declined but was still high in the peripheral region, consistent with a center-to-periphery retinal development pattern (Figure 2O). Meanwhile, Atoh7 expression expanded to the periphery, and reached peak levels (Figure 2O’). Although we observed a subset of cells co-expressing Atoh7 and Foxp1 at E14.5, most Atoh7+ cells no longer expressed Foxp1 (Figure 2R-2R’’), consistent with downregulation of primary RPC genes in neurogenic RPCs (10). At E16.5, both Foxp1 and Atoh7 expression were dramatically reduced in the central retina but remained high in the peripheral region (Figure 2P-2P’’). Cells co-expressing Foxp1 and Atoh7 were almost absent in the central region (Figure 2S-2S’’) but were detected at the periphery (Figure 2T-2T’’). Overall, we show that Foxp1 and Atoh7 have overlapping spatiotemporal expression patterns, but Foxp1 is more widely expressed than Atoh7, and its expression precedes Atoh7. More importantly, the colocalization of Foxp1 and Atoh7 supports the possibility that Foxp1 promotes the transition of RPCs to a neurogenic state by activating Atoh7.

### Atoh7 SE is required for the increase in Atoh7+ and Brn3b+ cells induced by Foxp1

To determine how Foxp1 regulates *Atoh7* expression, we assessed Foxp1 binding at the *Atoh7* locus. Although no Foxp1 peaks were detected at the *Atoh7* promoter or the primary enhancer (PE) region from Foxp1 CUT&RUN analysis, we identified a Foxp1 consensus peak at the remote shadow enhancer (SE) element, specifically within an evolutionarily conserved noncoding region which is open in ATAC-seq profiles and coincident with the active enhancer mark H3K27Ac (Figure 3A)(18, 20). These findings indicate that *Foxp1* directly activates *Atoh7* expression by binding to its SE element.

**Figure 3:**
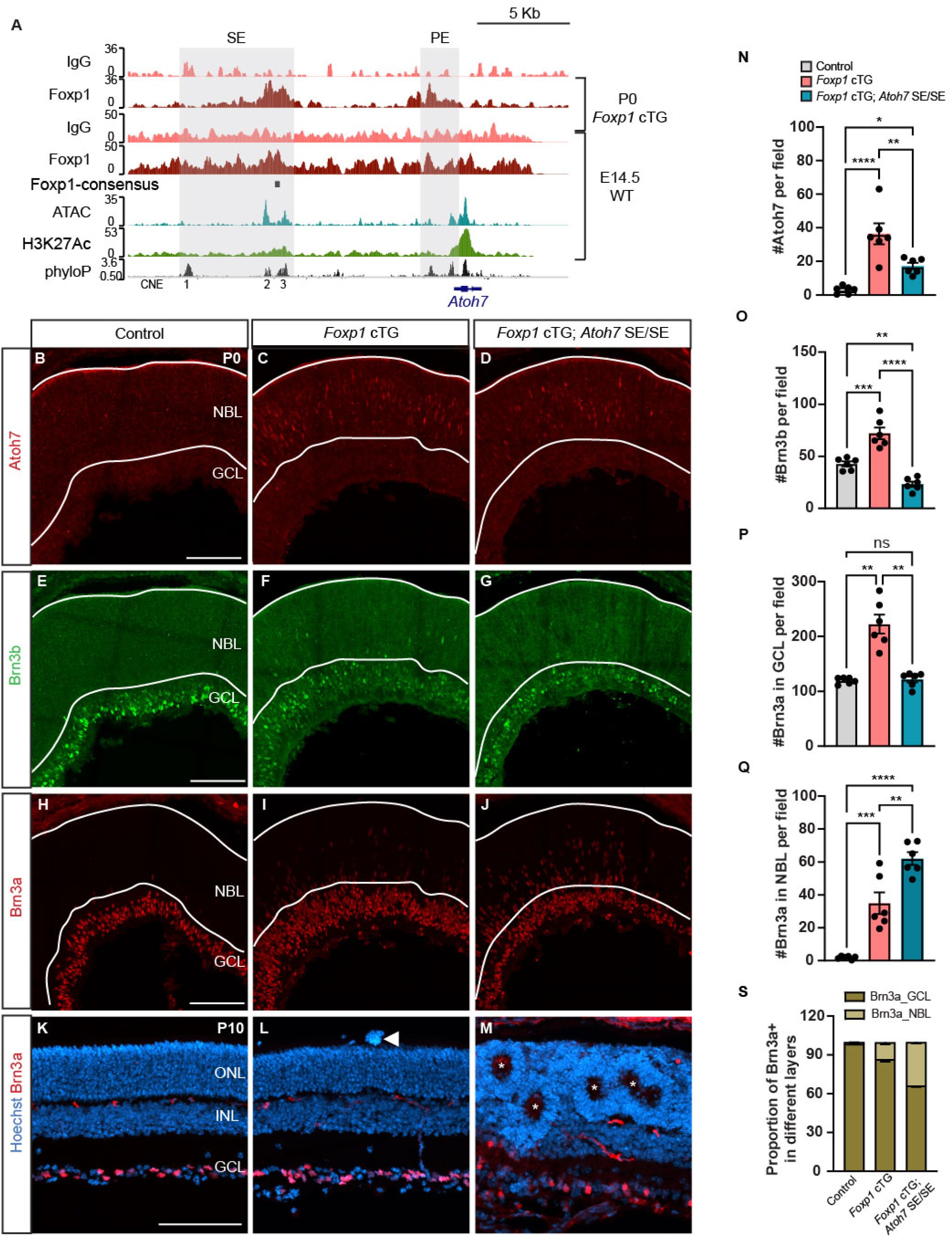
The Atoh7 SE is required for Foxp1-mediated Atoh7 activation and RGC genesis. (A) Foxp1 and IgG CUT&RUN genome tracks near the *Atoh7* locus in P0 *Foxp1* cTG and E14.5 WT retina. H3K27Ac ChIP-seq data and ATAC-seq data are from (68) and phyloP conservation tracks are from the UCSC genome browser (69, 70). The primary enhancer (PE), shadow enhancer (SE), and conserved non-coding elements (CNEs) of *Atoh7* are highlighted. (B-J) Representative images showing (B-D) Atoh7, (E-G) Brn3b, and (H-J) Brn3a in the central region of P0 (B, E, H) control (*Foxp1*^tg/+^), (C, F, I) *Foxp1* cTG and (D, G, J) *Foxp1* cTG; *Atoh7* SE/SE retina. (N-O) Quantification of the number of (N) Atoh7+ and (O) Brn3b+ cells per field within the central dorsal retina in different genotypes (n=6 retinas for each group). (N) *p=0.0439, **p=0.0071, ****p<0.0001 by one-way ANOVA with Tukey’s multiple comparisons. (O) **p=0.0061, ***p=0.0002, ****p<0.0001 by one-way ANOVA with Tukey’s multiple comparisons. (P-Q) Quantification of the number of Brn3a+ cells per field within the central dorsal retina in the (P) GCL and (Q) NBL of different genotypes (n=6 retinas for each group). (P) **p<0.01, ns: not significant by one-way ANOVA with Dunnett’s T3 multiple comparisons; (Q) **p=0.0017, ***p=0.0003, ****p<0.0001 by one-way ANOVA with Tukey’s multiple comparisons. (S) The proportion of Brn3a+ cells in the GCL and NBL in different genotypes. (K-M) Representative images showing Brn3a in the cross-section of P10 (K) control, (L) *Foxp1* cTG and (M) *Foxp1* cTG; *Atoh7* SE/SE retina. Arrowhead indicates abnormal subretinal structure. Asterisk marks rosette structure. Scale bar: 100 μm in all images. NBL: neuroblastic layer; GCL: ganglion cell layer; ONL: outer nuclear layer; INL, inner nuclear layer. Graphs represent mean ± SEM.

The SE element plays an important quantitative role by primarily regulating levels but not timing of *Atoh7* expression (18). To test whether the SE element is required for the increase in Atoh7+ cells induced by *Foxp1* overexpression, we combined the *Atoh7* SE deletion with *Foxp1* overexpression by crossing *Six3*-Cre, *Foxp1*^tg/+^ mice with *Atoh7* SE deletion mice (Atoh7 SE/SE)(18). Consistent with the previous report (19), Atoh7+ cells were almost absent in the central region of the control retina at P0 (Figure 3B). In contrast, the *Foxp1* cTG retina showed a substantial increase in Atoh7+ cells at P0 (Figure 3C, 3N), similarly to what we observed at E16.5 (Figure 2I, 2K). Deletion of the *Atoh7* SE in *Foxp1* cTG significantly reduced Atoh7+ cells compared to *Foxp1*-cTG alone, indicating that *Atoh7* SE depletion could reverse the increase in Atoh7+ cells caused by Foxp1 overexpression in RPCs, though the number of Atoh7+ cells were slightly higher than controls (Figure 3D, 3N), which might be caused by indirect regulation from *Foxp1*.

Atoh7 is necessary for the RGC cell fate and survival, and its dosage is correlated to RGC genesis (18, 21-23). We reasoned that RGC phenotypes would correspondingly change with the different Atoh7 levels present in our mice. We first focused on Pou4f2/Brn3b, which is a downstream target of Atoh7 and one of the earliest RGC markers (21). Consistent with the dramatic increase of Atoh7+ cells in *Foxp1* cTG, we observed a corresponding increase in the number of Brn3b+ cells (Figure 3E, 3F, 3O). Deleting *Atoh7* SE in *Foxp1* cTG mice significantly reduced the number of Brn3b+ cells induced by *Foxp1* elevation (Figure 3G, 3O). Notably, *Foxp1*-cTG; Atoh7 SE/SE retinas had fewer Brn3b+ cells than control retinas. Thus, *Atoh7* SE is required for the increase in Atoh7+ cells and Brn3b+ RGCs induced by Foxp1 elevation, supporting the conclusion that Foxp1 regulates Atoh7 expression and thus RGC production through the *Atoh7* SE element.

### Deleting the Atoh7 SE in *Foxp1*-cTG induced misplaced RGCs

To verify that Foxp1 regulates the production of fully differentiated RGCs through the *Atoh7* SE element, we extended our analysis to other RGC markers, including another known Atoh7 target, Pou4f1/Brn3a (21, 22). Brn3a+ cells dramatically increased in the *Foxp1* cTG retina at P0, leading to a thickened ganglion cell layer (GCL; Figure 3H, 3I, 3P), as we previously reported (5). After deleting *Atoh7* SE in *Foxp1*-cTG, we observed two distinct effects. Brn3a+ cells in the GCL were significantly reduced after deleting *Atoh7* SE in *Foxp1*-cTG, similar to what we observed for Brn3b (Figure 3J, 3P). However, we noticed a substantial increase in Brn3a+ cells in the neuroblastic layer (NBL) of *Foxp1* cTG; Atoh7 SE/SE (Figure 3J, 3Q), which we had not observed for Brn3b. We found the proportion of Brn3a+ cells in the NBL of *Foxp1*-cTG; *Atoh7* SE/SE retina was substantially higher than that of *Foxp1*-cTG only (Figure 3I, 3Q, 3S). We used additional RGC markers to characterize the misplaced Brn3a+ cells in *Foxp1*-cTG and *Foxp1*-cTG; *Atoh7* SE/SE and found that they express RBPMS, Isl1/2, and HuC/D (Figure S1). These data suggest that these Brn3a+ cells located in the NBL likely represent differentiated RGCs, but lack expression of Brn3b, a key factor in normal RGCs.

It has been reported that ectopically differentiated RGCs affect mature retinal lamination, resulting in the formation of rosette structures (24). Examining retinal lamination at P10, we had previously shown that most sections from the *Foxp1*-cTG retina have normal lamination, but some exhibited abnormal structures with aberrant cells protruding into the subretinal space (Figure 3L)(5). Notably, we observed numerous rosette structures in the *Foxp1*-cTG; *Atoh7* SE/SE retina, which had more misplaced RGCs at P0 (Figure 3M). Altogether, these results indicate that deleting *Atoh7* SE in *Foxp1*-cTG mice induced RGC-like cells that were misplaced in the NBL, affecting retinal lamination and leading to the formation of rosettes.

### Foxp1 promotes Neurog2 expression independently of Atoh7

We then asked how Foxp1 is promoting neurogenic progression and differentiation of these misplaced RGC-like cells when *Atoh7* levels are reduced by deletion of its SE. Since *Foxp1*-cTG *Atoh7* SE/SE had fewer Atoh7+ cells but more Brn3a+ RGCs in the NBL when compared to *Foxp1*-cTG alone, we concluded that other factors might be involved in Brn3a+ RGC genesis. To identify these factors, we performed bulk RNA-seq of P0 *Atoh7* SE/SE, *Foxp1* cTG, *Foxp1* cTG; *Atoh7* SE/SE and no Cre (control) retinas. We identified a total of 5,013 genes differentially expressed when each genotype was compared to the control. To understand how RGC genes were regulated in these conditions, we selected 69 transcription factors that have been identified as regulators of RGC development (25) and found that 40 of these were significantly changed in at least one comparison. To reveal the potential genetic regulatory relationship of these RGC-related DEGs, we grouped them into 6 clusters based on their expression pattern across different genotypes using the hierarchical clustering from the pheatmap package (Figure 4A).

**Figure 4:**
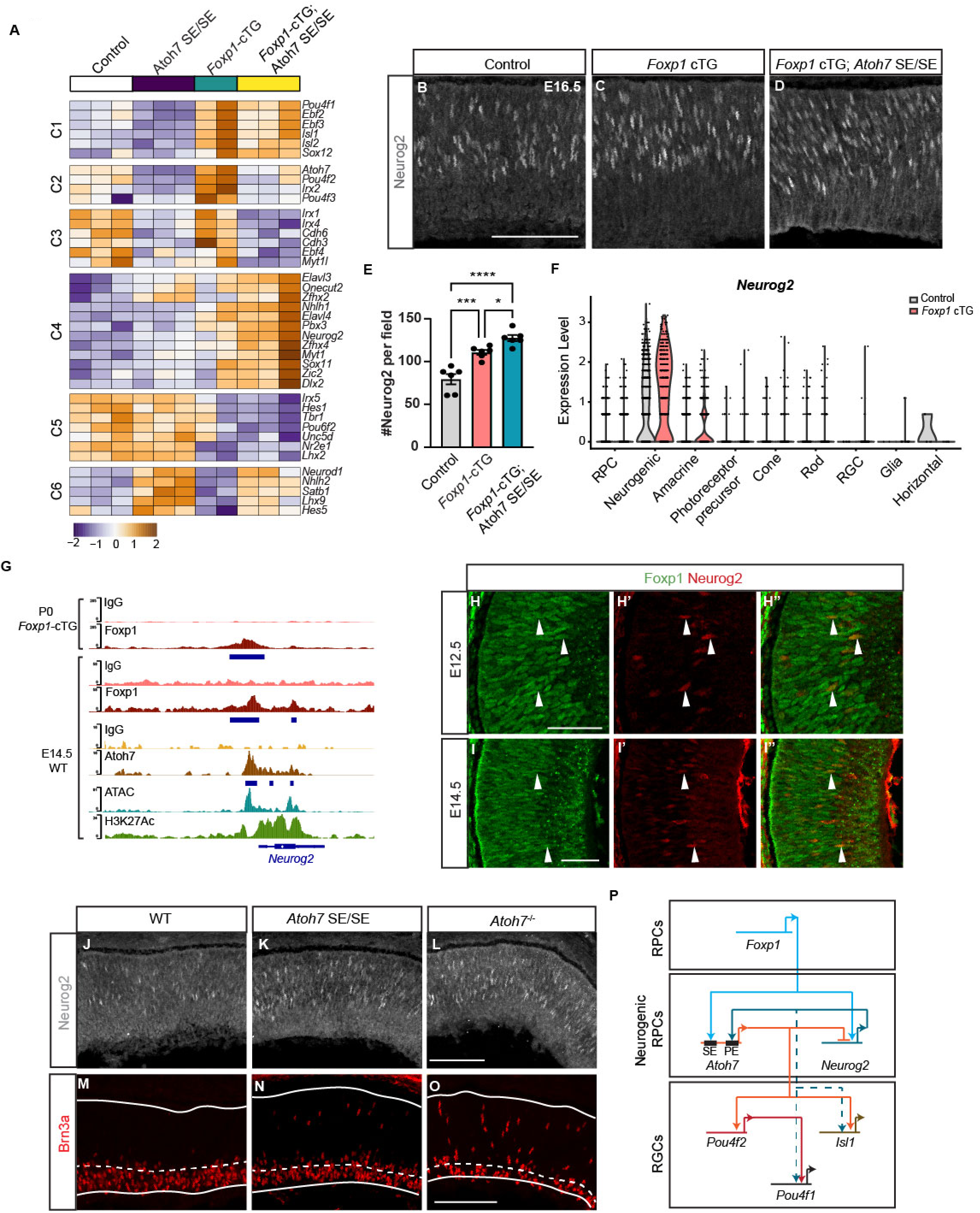
Foxp1 directly activates Neurog2 independent of Atoh7. (A) Heatmap of six clusters of RGC-associated transcription factors differentially expressed in *Atoh7* SE/SE, *Foxp1* cTG or *Foxp1* cTG; *Atoh7* SE/SE retina. (B-D) Representative images of Neurog2 in the central retina of E16.5 (B) control, (C) *Foxp1* cTG and (D) *Foxp1* cTG *Atoh7* SE/SE. (E) Quantification of Neurog2+ cells per field within the central dorsal retina in different genotypes at E16.5. *p<0.05, ***p<0.001, ****p<0.0001 by one-way ANOVA with Tukey’s multiple comparisons. (F) Violin plot of *Neurog2* expression by cell type from P0 *Foxp1* cTG and control scRNA-seq. (G) Foxp1 and IgG CUT&RUN genome tracks at *Neurog2* locus in P0 *Foxp1* cTG and E14.5 WT retina. Atoh7 CUT&Tag data from (22). H3K27Ac and ATAC-seq are from (68). (H, I) Single-z plane of confocal images showing Foxp1 and Neurog2 in the central region of (H) E12.5 and (I) E14.5 mouse retina. Arrowheads indicate Foxp1 and Neurog2 double positive cells. Scale bar: 50 μm. (J-L) Neurog2 immunofluorescence in E16.5 (J) WT, (K) *Atoh7* SE/SE mutant, and (L) *Atoh7*^-/-^mutant retina. (M-O) Representative images of Brn3a in E16.5 (M) WT, (N) *Atoh7* SE/SE mutant, and (O) *Atoh7*^-/-^mutant retina. Scale bar: 100 μm. (P) Gene regulatory network illustrating how Foxp1 activates Atoh7 and Neurog2, and the subsequence cross-regulation between these two bHLH factors drives distinct RGC genesis pathways.

This analysis confirmed that *Atoh7* was significantly downregulated in the *Atoh7* SE deletion retina (Figure 4A), as previously shown (18). We also verified that *Atoh7* was elevated in *Foxp1* cTG, which was reversed in *Foxp1* cTG; *Atoh7* SE/SE retina (Figure 4A), consistent with Foxp1 acting through the SE to drive *Atoh7* expression. We found that *Pou4f2 (Brn3b)* expression closely followed *Atoh7 expression* across different genotypes (C2, Figure 4A), while *Pou4f1 (Brn3a)* and *Isl1* (21, 22) clustered differently, despite being *Atoh7* downstream targets (21, 22). *Pou4f1* and *Isl1* were upregulated in *Foxp1* cTG retina and maintained high expression after deleting Atoh7 SE in *Foxp1* cTG mice (C1, Figure 4A), strongly indicating that they are also regulated by factors other than Atoh7.

From this analysis we identified neurogenic bHLH factor *Neurog2* as a likely driver of RGC gene expression. *Neurog2* was significantly upregulated in retinas from both *Atoh7* SE/SE and *Foxp1* cTG and was further increased in *Foxp1* cTG with *Atoh7* SE deletion (C4, Figure 4A-4F). Prior studies have identified a cross-regulatory relationship between Neurog2 and Atoh7: *Neurog2* directly activates Atoh7 expression through its primary enhancer (PE) whereas *Atoh7* represses Neurog2 as cells enter the nascent RGC state, with *Atoh7* loss resulting in *Neurog2* de-repression (26-30). We therefore expected *Neurog2* de-repression and upregulation in the *Atoh7* SE/SE retina as observed. However, we unexpectedly found Neurog2 elevated in the *Foxp1* cTG retina even though *Atoh7* was increased, indicating that *Foxp1* could independently activate *Neurog2* (Figure 4A-4F). Indeed, we noted a Foxp1 consensus peak at the promoter region of *Neurog2* (Figure 4G) and found co-expression of endogenous Foxp1 and Neurog2 in the embryonic mouse retina (Figure 4H-4I). These results suggest that *Neurog2* is directly activated by *Foxp1* and repressed by *Atoh7,* resulting in elevated *Neurog2* levels in *Foxp1* cTG retinas after deletion of *Atoh7* SE (Figure 4G). Our findings indicate that *Foxp1* promotes neurogenic progression by activating both *Atoh7* and *Neurog2,* while these bHLH factors cross-regulate each other to coordinate early retinal neurogenesis (Figure 4P).

We noted that the upregulation of *Neurog2* in both *Foxp1* cTG and *Foxp1* cTG; *Atoh7* SE/SE retina was correlated with increased Brn3a+ cells misplaced in the NBL (Figure 3P,3Q). To further test whether *Neurog2* level is associated with the misplaced RGCs in the NBL, we examined *Atoh7* SE/SE and *Atoh7*^-/-^mice (31), in which Atoh7 is low or absent while Neurog2 is high (Figure 4J-4L). By performing Brn3a immunostaining at E16.5, we found misplaced Brn3a+ cells in the NBL of both *Atoh7* SE/SE and *Atoh7*^-/-^ mice (Figure 4M-4O). Consistent with our model, Neurog2 was visibly elevated in *Atoh7*^-/-^ retina compared to *Atoh7* SE/SE with a striking increase in the proportion of Brn3a+ cells displaced into the NBL. Together these findings imply that although Neurog2 can partially compensate for the absence of Atoh7 to drive RGC genesis, the resulting RGCs exhibit abnormalities in both transcriptomic profiles and spatial distribution. Overall, we conclude that Foxp1 normally initiates the transition of RPCs to the neurogenic state by activating both Atoh7 and Neurog2, with cross-regulation between these factors required to coordinate the normal differentiation of mature RGCs.

### Foxp1 is sequentially marked by H2AK119ub1 and H3K27me3

Having established that Foxp1 directly activates proneural genes, we next asked how this regulatory activity is restricted over developmental time by examining the chromatin changes and cis-regulatory elements that establish repression of *Foxp1* in RPCs. Previously, we found that the repression of *Foxp1* in RPCs during development is associated with de novo establishment of the PRC2 complex-mediated repressive histone modification H3K27me3 at its genomic locus (5), but how this is established and the genomic regions critical for Foxp1 repression are undefined. To better understand how H3K27me3 accumulates, we performed CUT&RUN for H3K27me3 at multiple developmental stages and found that H3K27me3 signals were absent at the *Foxp1* locus at E14.5 when Foxp1 levels are high, first detectable at E15.5, slightly increased at E16.5, and robustly deposited at P0 when Foxp1 is fully repressed in RPCs (Figure 5A). It is known that H3K27me3 tends to colocalize with PRC1 complex-mediated H2AK119ub1 to maintain transcriptional repression in different contexts (32-34). Since the genomic distribution of H2AK119ub1 in developing retina was undefined, we performed H2AK119ub1 CUT&RUN at the same developmental points to characterize H2AK119ub1 deposition at *Foxp1*. We found that H2AK119ub1 colocalized with H3K27me3 at the *Foxp1* locus and gradually increased over time (Figure 5A). Notably, weak H2AK119ub1 peaks were already present across *Foxp1* by E14.5, a time point when H3K27me3 was absent, suggesting that H2AK119ub1 precedes H3K27me3 deposition at the *Foxp1* locus (Figure 5A).

**Figure 5:**
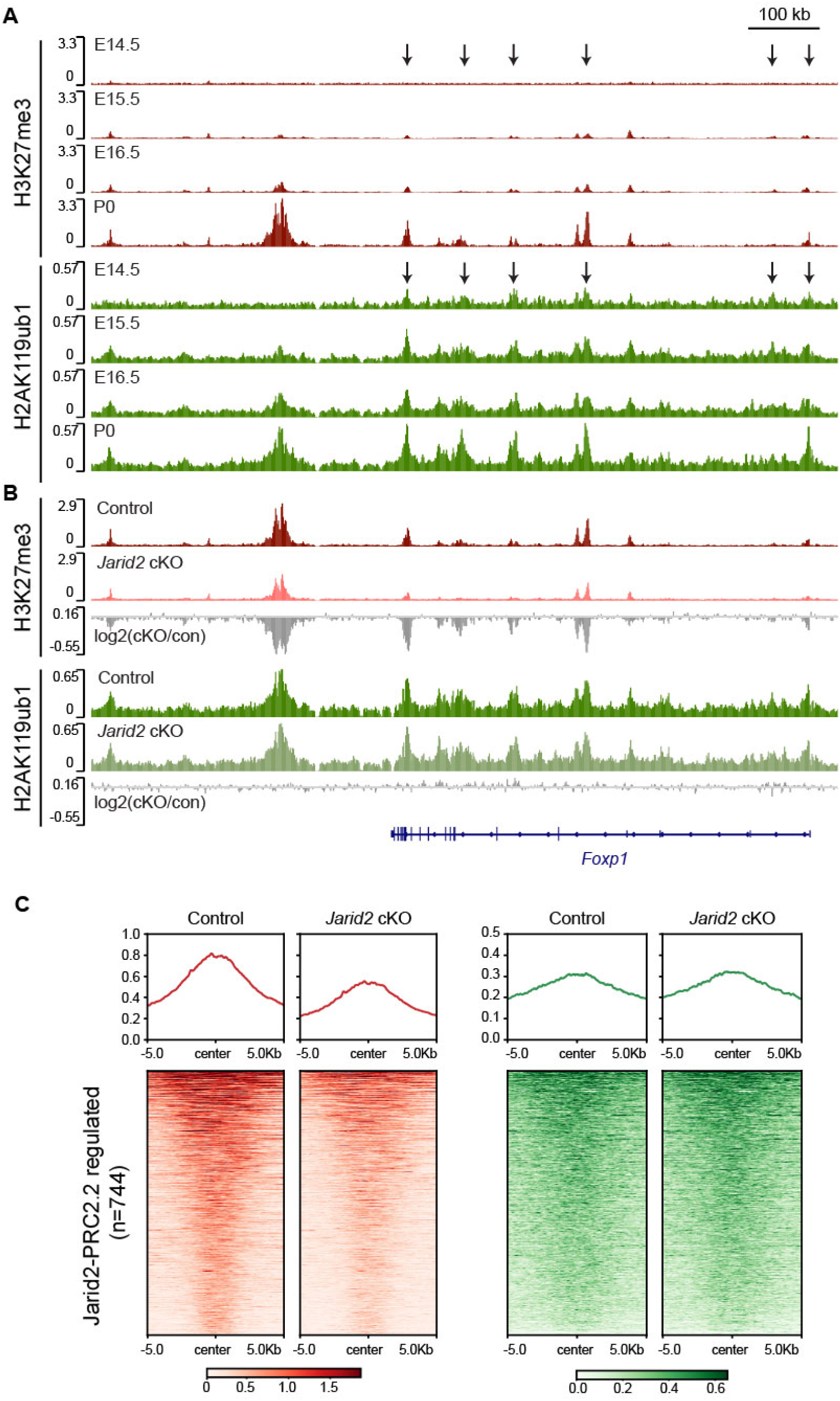
H2AK119ub1 and H3K27me3 are sequentially deposited at the *Foxp1* locus. (A) CUT&RUN tracks of H3K27me3 and H2AK119ub1 at the *Foxp1* locus from E14.5, E15.5, E16.5 and P0 WT mouse retina. Arrows indicate H2AK119ub1 peaks present at E14.5. (B) CUT&RUN tracks of H3K27me3 and H2AK119ub1 within *Foxp1* genomic locus from *Jarid2* control and cKO retina. (C) Heatmap and profile plot showing average H3K27me3 and H2AK119ub1 signal across regions with reduced H3K27me3 in *Jarid2* cKO retina identified in (5). H3K27me3 datasets in (B-C) are from (5).

Prior studies suggest reciprocal models for Polycomb recruitment: H3K27me3 deposited by PRC2 can recruit canonical PRC1 through its CBX subunits, whereas H2AK119ub1 deposited by variant PRC1 can bind JARID2 and recruit PRC2, promoting subsequent H3K27me3 deposition (35), so we sought to distinguish these two possibilities. We previously showed that the PRC2.2 subunit *Jarid2* is important for H3K27me3 deposition at the *Foxp1* locus (5), so we first investigated whether Jarid2 regulates the deposition of H2AK119ub1 by performing CUT&RUN in E18.5 *Rax*-Cre^ERT2^; *Jarid2*-cKO and littermate control mice. While H3K27me3 at *Foxp1* was significantly reduced, we did not observe significant changes in H2AK119ub1 at *Foxp1* after depleting *Jarid2* in RPCs (Figure 5B).

Neither did we find changes in H2AK119ub1 at other Jarid2-PRC2.2 regulated regions in the *Jarid2*-cKO retina (Figure 5C). Overall, these data demonstrate that recruitment of PRC1 to the *Foxp1* locus is independent of Jarid2/PRC2.2-mediated deposition of H3K27me3, suggesting that sites are pre-marked with H2AK119ub1 to promote recruitment of Jarid2/PRC2.2 and deposition of H3K27me3 resulting in progressive repression of Foxp1.

We next sought to investigate the mechanisms by which polycomb repressive complexes are recruited. The Casz1 protein, especially the short splice variant Casz1v2, has been reported to interact with PRC1 core component, Ring1b, in the mouse retina (36, 37). Moreover, Foxp1 expression declines in RPCs when Casz1 begins to be expressed (Figure S2D) (5, 14). This raises the possibility that Casz1 recruits PRC1 complex to repress Foxp1 via interaction with Ring1b. To test this, we performed Casz1 CUT&RUN in the E18.5 and P0 mouse retina, where Casz1 is highly expressed, and identified Casz1 peaks on the *Foxp1* locus (Figure S2A). Interestingly, the Casz1 binding site on Foxp1 as well as other regions along the Foxp1 locus are enriched for Ring1b (Figure S2A and S2C). We then examined whether *Casz1* is required for the deposition of H2AK119ub1 and H3K27me3 on *Foxp1*, using *Chx10-Cre*; *Casz1*^fl/fl^ to conditionally delete *Casz1* in RPCs. However, we found the enrichment of H2AK119ub1 and H3K27me3 on *Foxp1* did not change significantly (Figure S2B). We further tested whether Casz1 regulates Foxp1 expression by electroporating E14.5 mouse retinal explants with GFP or Casz1v2-IRES-GFP. After 48 hours, we sorted GFP+ cells and extracted total RNA for RT-qPCR (Figure S2D). We found that overexpression Casz1 did not alter Foxp1 expression level (Figure S2E). We conclude that Casz1 is not required for the recruitment of polycomb repressive complexes to *Foxp1*, even though it is associated with Ring1b, indicating that other factors may be involved.

### A distal conserved region functions as a silencer of *Foxp1*

To better understand how *Foxp1* is gradually repressed in RPCs, we used published scATAC-seq datasets from the developing mouse retina to examine chromatin accessibility around *Foxp1* (11). We identified a distal intergenic non-coding region located approximately 150 kilobase (kb) downstream of *Foxp1* that was closed at E14 and gradually opened from E16 in RPCs, coincidence with declining expression of *Foxp1* (Figure 6B). Surprisingly, we noticed that the region was specifically accessible in RPCs at P0 but not in other cell types, including RGCs, in which Foxp1 is expressed. This distal region shows high evolutionary conservation, suggesting a crucial developmental function; hereafter, it is referred to as the DisCo (<u>Dis</u>tal <u>Co</u>nserved) region (Figure 6B). Though linearly distant from *Foxp1*,analysis of published Hi-C data from P0 mouse retina revealed that the DisCo region and *Foxp1* reside within the same topologically associating domain (TAD), suggesting physical proximity (Figure 6A and Figure S3). We examined histone modifications around the DisCo region at P0 to predict its function and found the active enhancer mark H3K27Ac was absent, but repressive marks H3K27me3 and H2AK119ub1 were strongly and broadly enriched (Figure 6B). Ring1b is present in a narrow peak centered in the DisCo region, supporting this as the site for nucleation of these repressive marks.

**Figure 6:**
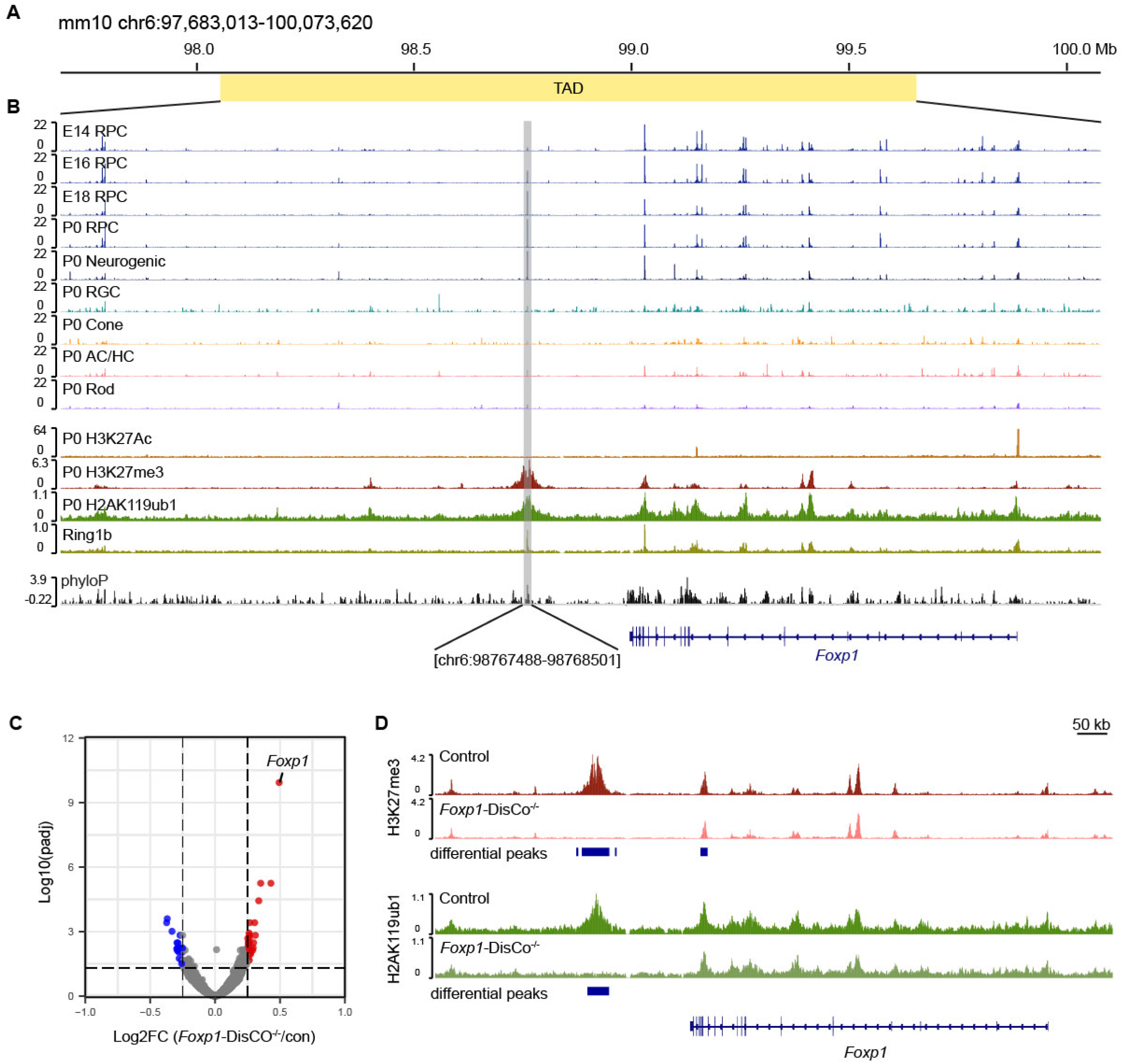
A distal H3K27me3-enriched region functions as a silencer of *Foxp1*. (A) A genomic region spanning *Foxp1* (chr6: 97,683,013-100,073,620; mm10). The yellow region indicates the predicted TAD in which *Foxp1* resides, from previously published Hi-C data in mouse retina (71). (B) Genome tracks across the TAD, from top to bottom: (1) scATAC-seq tracks showing chromatin accessibility in RPCs across development stages and in different retinal cell types at P0 from developing mouse retina (11); (2) tracks of epigenetic markers H3K27Ac, H3K27me3, H2AK119ub1 and Ring1b in E18.5 or P0 mouse retina; (3) phyloP conservation track for 60 species. Highlighted region is the putative silencer element of *Foxp1*. Genomic coordinates indicate the *Foxp1* distal conserved region deleted in DisCo knockout mice. P0 H3K27Ac ChIP-seq data is from (68) and phyloP conservation tracks are from the UCSC genome browser (69, 70). (C) Volcano plot of differentially expressed genes in P0 *Foxp1*-DisCo^-/-^mice (n = 3 for each genotype; adjusted *p* < 0.05). Red: significantly upregulated genes; blue: significantly downregulated genes. (D) CUT&RUN tracks of H3K27me3 and H2AK119ub1 in P0 *Foxp1*-DisCo^-/-^and littermate WT mice around the *Foxp1* loci. Differential peaks were identified by DESeq2 analysis.

H3K27me3-rich genomic regions have been reported as a marker of transcriptional silencers (38-40). This is consistent with our observation that the accessibility of the DisCo region was negatively associated with *Foxp1* expression levels. We hypothesized that the DisCo region acts as a silencer element to gradually repress *Foxp1* as RPCs mature.

To test this hypothesis, we generated a mouse line with constitutive deletion of the DisCo region. By performing bulk RNA-seq analysis at P0, we found that *Foxp1* was the most significantly upregulated gene among the total 177 differentially expressed genes in *Foxp1*-DisCo^-/-^ retina (Figure 6C). This result supports the conclusion that the DisCo region acts as a silencer of *Foxp1*. As the DisCo region is strongly associated with broad H3K27me3 and H2AK119ub1 signals, we tested whether it is required for the deposition of these two repressive histone marks by performing CUT&RUN in P0 *Foxp1*-DisCo^-/-^and littermate controls. We found that H3K27me3 and H2AK119ub1 failed to deposit around the deleted region in *Foxp1*-DisCo^-/-^ retina while H3K27me3 was also significantly reduced at neighboring peaks. Overall, these results demonstrate that the DisCo element is required for the local deposition and spreading of repressive H3K27me3 and H2AK119ub1 marks, thus playing a key role in the repression of Foxp1 in RPCs.

## Discussion

Here, we show that temporal factor *Foxp1* plays dual roles in maintaining the early RPC state and promoting the transition to the early neurogenic state. We report that *Foxp1* directly activates early RPC genes while repressing late temporal RPC factors. We also establish a direct regulatory relationship between *Foxp1* and the early proneural factors *Atoh7* and *Neurog2*, and demonstrate that disrupting this relationship leads to abnormal RGC genesis, while increasing levels of Foxp1 expression directly drives increased RGC genesis. We showed that temporal restriction of this early program occurs via progressive silencing through sequential deposition of repressive histone modifications

H2AK119ub1 and H3K27me3 at the *Foxp1* locus, and we specifically identify a conserved distal H3K27me3-enriched region that functions as a silencer of *Foxp1*. Overall, these findings reveal a central role for Foxp1 in coordinating early temporal patterning, neurogenic progression, and the generation of early born retinal neurons.

We established that Foxp1 functions as both a transcriptional activator and repressor in RPCs. Gene regulatory networks (GRNs) constructed by integrating scRNA-seq and scATAC-seq data from developing mouse retina predict that temporal transcription factors preferentially activate factors within their own network and repress those in the opposing network (11). We provide direct evidence for this, showing that Foxp1 directly activates other early factors (e.g., *Ndufa4*, *Col9a1*) and represses late temporal factors, including the well-established *Casz1* and *Nfib* (10, 14). Although all three Nfi family members (Nfia, Nfib, and Nfix) are enriched in late RPCs and determine late RPC identity (10), we found only *Nfib* was significantly changed in *Foxp1* manipulated retinas, and *Foxp1* peaks were detected exclusively at the *Nfib* locus, suggesting specific regulation of *Nfib* by Foxp1 in the developing mouse retina. The GRNs suggest mutual repression between factors in early and late-RPCs. Such mutual repression between early and late-stage transcription factors has been identified in other developing CNS regions, including the developing spinal cord (41). We observed that the late factor Casz1 binds to *Foxp1* locus, including at the silencer-like DisCo region. However, overexpressing *Casz1* in E14.5 mouse retina did not affect *Foxp1* expression, suggesting Casz1 alone does not repress *Foxp1*. While it is unknown whether Nfi factors repress *Foxp1* in the developing retina, we analyzed published Nfia/b/x ChIP-seq data from P2 mouse retina (11) and identified an Nfia/b/x peak overlapping with Casz1 in the DisCo region of *Foxp1* (not shown). Thus, it is possible that Nfi factors, potentially in cooperation with Casz1, regulate *Foxp1* expression. Overall, our findings define how Foxp1 activates early factors and represses late temporal factors to sustain early temporal competence in RPCs.

While Foxp1 is primarily known as a transcriptional repressor, prior evidence suggests it can also function as an activator in certain contexts, including in the developing mouse cortex (17, 42-45). However, the underlying mechanisms remain incompletely understood. One possibility is that Foxp1 interacts with distinct context-specific co-factors to exert its functions. Foxp1 is known to interact with the co-repressor CtBP1 and repressive NuRD complexes (46), as well as other neurodevelopmental transcription factors, including NR2F1/2 and Ikaros (47, 48). Notably, NR2F1/2 and Ikaros are co-expressed with Foxp1 in RPCs during the same developmental window (12, 49). It is yet unclear whether Foxp1 interacts with these or other factors in the developing mouse retina to perform its dual functions.

An unresolved question in development has been how primary RPCs progress to the neurogenic state to initiate neuron differentiation (50). We provide evidence that Foxp1 directly activates the proneural genes *Atoh7* and *Neurog2* via binding to the *Atoh7* SE and *Neurog2* promoter, thus linking Foxp1 expressed in early RPCs with neurogenic progression. We found that elevating *Foxp1* increases the number of Atoh7+ cells and differentiated RGCs, and this was dependent on the *Atoh7* SE. Consistent with this, two independent studies demonstrated that the *Atoh7* SE region, where Foxp1 binds, is required to modulate both *Atoh7* expression level and RGC genesis (18, 20).

It has been shown that *Neurog2* acts at the leading edge of retinal neurogenesis and directly activates *Atoh7*, whereas *Atoh7* inhibits expression of *Neurog2* (26-29). While *Foxp1* activates both *Atoh7* and *Neurog2*, only *Atoh7* was downregulated in *Foxp1* cKO retina while *Neurog2* remained unchanged, potentially resulting from de-repression due to reduced *Atoh7*. Based on this cross-regulatory relationship, increased Neurog2 expression might contribute to the remaining Atoh7+ cells we observed after deleting *Atoh7* SE in *Foxp1* cTG retina. Increased Neurog2 was also associated with elevated expression of a subset of differentiated RGC genes including Pou4f1 but not Pou4f2. Consistent with this, in *Foxp1* cTG retina after *Atoh7* SE deletion, Pou4f1/Brn3a+ RGCs were increased, with misplaced Brn3a+ RGCs evident in the NBL while Atoh7+ cells and Pou4f2/Brn3b+ RGCs were decreased relative to *Foxp1* cTG alone. We also observed misplaced Brn3a+ RGCs in *Atoh7*^-/-^ retinas at embryonic stages. These observations suggest an Atoh7-independent mechanism for generating Brn3a+ RGCs that are misplaced and partially differentiate but lack expression of Pou4f2/Brn3b. This model is supported by the observation that *Atoh7*^-/-^ mice, with apoptosis inhibited, achieve near-normal levels of Pou4f1/Brn3a RGCs, while Pou4f2/Brn3b RGCs are reduced (51). Thus, while Atoh7-independent mechanisms can generate RGCs, our findings show that their differentiation is incomplete, potentially explaining why they ultimately die in the *Atoh7*^-/-^ model.

We argue that Neurog2 can act to promote this incomplete RGC genesis. Neurog2 and Atoh7 are both activated in neurogenic RPCs during retina development (27). Loss of *Neurog2* reduces Pou4f+ RGCs and downregulates RGC genes including *Pou4f1*, *Pou4f2*, *Onecut2*, *Onecut3*, and *Ebf3* (27, 52). Moreover, *NEUROG2* overexpression in iPSCs or human organoids induced the production of RGCs with low *POU4F2* but high *POU4F1*, distinct from those generated in control or by ATOH7 overexpression (53, 54). Together, these findings indicate that *Neurog2* promotes partial RGC differentiation, favoring Pou4f1/Brn3a-expressing RGCs that lack Pou4f2/Brn3b. We cannot exclude the possible involvement of other transcription factors known to promote RGC genesis, like *Onecut2* which has similar expression dynamics as Neurog2 across the genetic models we analyzed (55). Altogether, our findings support the conclusion that both Atoh7 and Neurog2 need to be maintained at a certain ratio to ensure normal RGC genesis, achieved partially through cross-regulation, with Foxp1 acting upstream to drive expression of both to promote the transition to the neurogenic state (Figure 4P).

We examined how sequential histone modifications and cis-regulatory elements at the Foxp1 locus restrict this early program as development proceeds. We previously showed that *Foxp1* must be repressed to ensure temporal progression of RPCs, and the timing of *Foxp1* decline in RPCs is correlated with the *de novo* accumulation of H3K27me3 mediated by the Jarid2/PRC2.2 complex (5). Here, we show that PRC1-mediated H2AK119ub1 is deposited at the *Foxp1* locus by E14.5, preceding H3K27me3. While traditional models suggest the recruitment of PRC1/H2AK119ub1 is dependent on PRC2/H3K27me3, there is strong evidence for an alternative model where H2AK119ub1 is deposited by PRC1 to target sites, then recognized and bound by Jarid2, leading to PRC2 recruitment and H3K27me3 deposition (56-58). The sequential deposition of H2AK119ub1 and H3K27me3 we observed at the *Foxp1* locus during mouse retina development strongly supports the alternative model of PRC1 recruiting PRC2.2. We show that depleting *Jarid2* reduces the enrichment of H3K27me3 without altering H2AK119ub1, indicating that Jarid2/PRC2.2-H3K27me3 acts downstream of PRC1-H2AK119ub1. Our findings are consistent with a report that in embryonic stem cells JARID2-sensitive genes are pre-marked by H2AK119ub1 and become newly repressed upon their differentiation to neural progenitor cells (59). The PRC1 complex also regulates temporal progression in other CNS regions. In the mouse neocortex, Ring1b and H3K27me3 progressively accumulate at the *Fezf2* locus, and depleting Ring1b or inactivating the PRC2 component Ezh2 de-represses *Fezf2* and increases the production of early-born neuronal subtypes (60), similar to Jarid2/PRC2.2 function in mouse retina development (5). The progressive accumulation of repressive PRC1/PRC2 marks at temporal factor loci in distinct CNS progenitors suggests a conserved mechanism mediating temporal identity transitions.

We addressed which cis-regulatory region of Foxp1 may be critical for its repression and identified a highly conserved silencer element for *Foxp1.* We observed striking negative correlation between *Foxp1* expression in RPCs and chromatin accessibly of this DisCo region, as well as the presence of broad H3K27me3 deposition, and further showed that deleting the DisCo region upregulates *Foxp1* expression in P0 mouse retina. Although the epigenetic signatures of silencers have not been as well defined as those of enhancers, several studies have suggested that H3K27me3-enriched accessible chromatin regions act as silencers in multiple contexts (40). The DisCo region is located in an intergenic region approximately 150kb downstream of the *Foxp1* promoter within the same TAD as *Foxp1*, but whether and how the DisCo region physically interacts with the *Foxp1* promoter remains unclear. In the DisCo^-/-^retina, the absence of H3K27me3 and H2AK119ub1 around the DisCo region and their reduction across the *Foxp1* locus indicate that the Polycomb complexes may mediate the silencer function of the DisCo region.

Overall, our findings identify Foxp1 as a mechanistic link between early progenitor identity and neurogenic progression, with distinct cis-regulatory elements mediating its control of neuronal production and temporal repression. This coordination suggests that temporal competence factors can shape neural diversity both by defining the fates available to progenitors and by engaging the transcriptional programs that initiate their differentiation. More broadly, the study provides a framework for investigating how developmental changes in progenitor identity are translated into temporally appropriate neuronal output across the CNS.

## Methods

### Animals

#### Mice

All animal procedures were approved by the University of Utah Institutional Animal Care and Use Committee (IACUC). C57BL/6 mice were purchased from the Jackson Laboratory. *Jarid2* conditional knockout (cKO) and *Foxp1* conditional transgenic (cTG) mice were generated as previously described (5). Briefly, *Jarid2* floxed mice were crossed with *Rax-CreER^T2^* (RRID: IMSR_JAX:025521). Tamoxifen (75 μg of 10 mg/ml per gram of body weight) was injected intraperitoneally to pregnant females at E10.5 to induce Cre recombinase activity. The day of vaginal plug detection was marked as E0.5. *Foxp1a* transgenic mice (MGI:5607369) were crossed with *Six3-Cre* mice to generate *Six3-Cre*; *Foxp1*^tg/+^ (*Foxp1*-cTG) and *Six3*-Cre; *Foxp1*^tg/tg^ mice (61, 62). To generate *Six3-Cre*; *Foxp1^fl^*^/fl^ mice, *Foxp1* floxed mice (RRID: IMSR_JAX:017699) were crossed with *Six3-Cre* mice (63). *Atoh7* SE deletion mutant mice were originally generated in Dr. Tom Glaser’s lab and were kindly provided by Dr. Joel B. Miesfeld (18). To delete *Atoh7* SE in *Foxp1*-cTG mice, *Atoh7* SE deletion mutant mice were crossed with *Six3*-Cre; *Foxp1*^tg/+^ mice. *Atoh7*^-/-^mice were from Dr. Tom Glaser’s lab (31). *Casz1* cKO mice were generated by crossing *Chx10*-Cre mice with *Casz1*^fl/fl^ (14), and were conducted according to the guidelines of the Canadian Council on Animal Care and the Animal Care and Veterinary Service at University of Ottawa using ethical protocols OHRI-2856 and OHRI-3949. *Foxp1* distal conserved region (*Foxp1*-DisCo) knockout mice were generated using CRISPR-Cas9 technology by the University of Utah Transgenic & Gene Targeting Mouse Core Facility. Single-guide RNAs (sgRNAs) targeting the DisCo region (chr6:98,767,488-98,768,501 in mm10) were designed and optimal pairs were selected based on predicted on-target efficacy and off-target scores. Cas9 protein and sgRNAs were delivered into single-cell mouse embryos by electroporation. F0 founder mice carrying the desired deletions were backcrossed to C57BL/6J to generate F1 offspring, and precise deletions were confirmed by Sanger sequencing. sgRNA sequences and primers for genotyping are listed in Table 1.

**Table 1.**
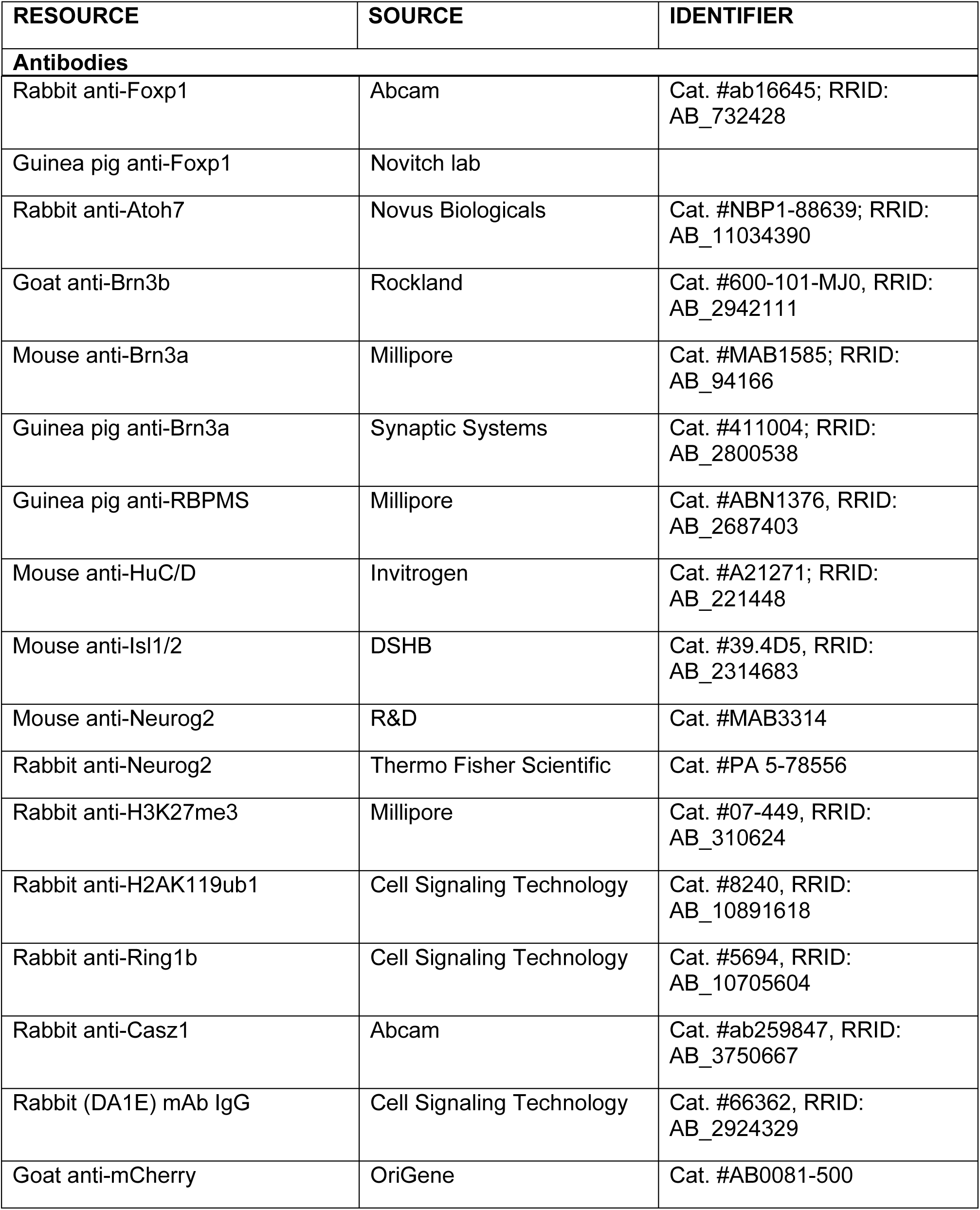
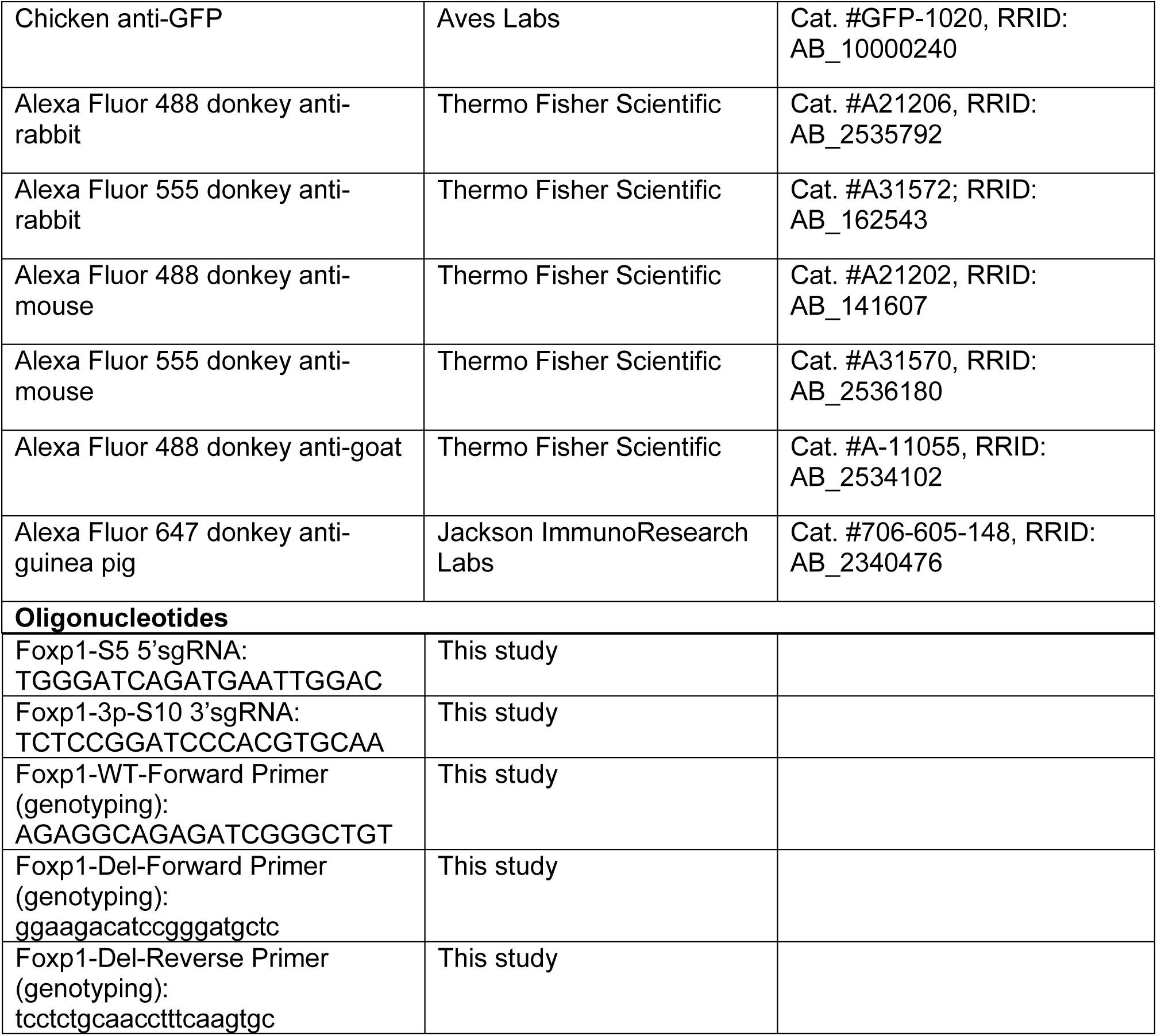
List of antibodies and oligonucleotides used in this study.

#### Zebrafish

All zebrafish work was performed in accordance with the Institutional Animal Care and Use Committee of the Medical College of Wisconsin. The transgenic line generated to overexpress mouse *Foxp1* utilized a bi-directional upstream activator sequence (UAS) to express both Foxp1 and a destabilized nuclear mGreenLantern (nmGrLPEST) simultaneously (nmGrLPEST:*UAS*:Foxp1*^mw802^*) and was driven by *vsx2*:Gal4-VP16*^mw39Tg^* (64). The human ATOH7 transgenic reporter line was generated by recombineering in Tol2 transposase sites and a destabilized nuclear mCherry (nmChPEST) fluorescent protein into human BAC (bacterial artificial chromosome) clone RP11-1114I5 (*HATOH7*:nmChPEST*^mw803^*). Each stable transgenic line was established as previously described (65). **Tissue processing**

For embryonic and P0 tissues, whole heads were collected and fixed in 4% PFA for 1 hour. For postnatal tissues, eyeballs were enucleated from euthanized mice, and the dorsal side was marked. Eyecups were prepared by fixing eyeballs in 4% PFA for 30 minutes after removing the cornea and lens. Tissues were washed three times in PBS for 20 minutes each and treated in 10% and 20% sucrose in PBS at 4°C. Samples were embedded in OCT compound (Tissue-Tek, Cat #27050) and stored at -80 °C. Coronal cryostat sections were made at 16 µm thickness.

### Immunohistochemistry, Confocal microscopy and Image quantification

Immunohistochemistry was performed as previously described (5). Briefly, retina sections were incubated in blocking buffer (0.2% triton-X, 10% BSA, 10% normal donkey serum in 0.01 M PBS) for 1 hour at room temperature. Sections were then incubated in primary antibody buffer (0.2% triton-X, 5% BSA in 0.01 M PBS) with desired antibodies overnight at 4 °C. After three washes in PBS, sections were incubated with secondary antibodies (5% BSA in PBS) for 2 hours at room temperature, washed, and mounted with Fluoromount-G mounting medium. All primary and secondary antibodies used in the paper are listed in Table1.

Confocal images were acquired on an inverted Nikon A1R Confocal Microscope. Images were collected with a 20X objective and a 3X digital zoom, yielding a 0.2 μm pixel resolution. Stacks through the Z plane were taken at 0.8 µm step distance, a total depth of about 16 μm, covering the entire retina thickness. Image acquisition settings were consistent across ages and genotypes. Confocal images shown are representative of at least 3 animals/6 retinas. For quantification, the number of retinal cells was quantified manually and blindly by averaging total positive cells within the central dorsal retinal region (∼300 μm long) from 4-5 sections per retina using Nikon Elements software.

Zebrafish whole mount and cryosection histochemistry was performed as previously described (65). Confocal images of whole mount retinas were taken on a Nikon Laser Scanning Confocal Processed using Fiji to incorporate a 4 μm z-stack of the retina. Cryosection images (4 μm) were captured on a Zeiss AxioImagerZ.2 and images processed using ZenBlue v3.5 software.

### Retinal explant electroporation and tissue collection

E14.5 eyes were dissected in Hank’s Balanced Salt Solution (HBSS). Lenses were not removed to prevent damage to the retina. The dissected retina was transferred to the electrode chamber containing 100 µl desired DNA plasmid solution (1 µg/µl). The pCAGIG and pCAG-Casz1v2 plasmids were kindly provided by Cayouette lab (13). The retina was oriented with the photoreceptor layer facing the negative electrode and then electroporated using five square wave pulses of 25 V for 50 ms with 950 ms intervals between pulses. Following electroporation, retinas were cultured on Millicell standing inserts in 24-well plates containing Neurobasal medium supplemented with 2% B27, 1% Pen-Strep, 1% L-Glutamine and 10% FBS for 48 hours.

### RT-qPCR

Retinas were dissociated using papain as described in the CUT&RUN experiment. Electroporated GFP+ cells were sorted directly into Qiagen Buffer RLT buffer using a BD FACSAria Cell Sorter. Only samples containing more than 1000+ sorted GFP cells were retained. RNA was extracted using the RNeasy Plus Mini Kit (Qiagen, Cat #74134). First-strand cDNA synthesis was performed using the ZymoScript RT PreMix Kit (Cat #R3012). Quantitative real-time PCR was performed using SYBR Select Master mix (Applied Biosciences, Cat #4472908) and run on the 7900 HT Fast Real-Time PCR system with QuantStudio 12K Flex software (Applied Biosystems) at the University of Utah Genomics Core. Beta Actin (Actb) was used as the housekeeping gene and the 2- ΔΔCT method was used to calculate relative fold changes in gene expression.

### CUT&RUN

#### Sample and Library Preparation

CUT&RUN experiments were performed as previously described (5). Fresh retinas were used in most experiments, except for *Casz1* cKO and littermate control retinas, which were cryopreserved in Neurobasal medium containing 10% DMSO and slowly frozen in an isopropanol-filled chiller at -80°C. Cryopreserved retinas were rapidly thawed at 37°C before CUT&RUN. Retinas were dissociated in a papain buffer (HBSS with 50 mM L-cysteine, 10 mM EDTA, 10 mM 2-Mercaptoethanol, and 10 mg/ml papain) with 1 mg/ml DNase I at 37°C for 10 min. Papain activity was quenched by adding ovomucoid solution. Cells were centrifuged at 300 g for 5 min. The pellet was resuspended in Neurobasal medium and filtered through a 40 μm Cell Strainer to get single-cell suspensions. CUT&RUN was performed following the manufacturer’s protocol (Cell Signaling Technology, Cat #86652). For each reaction, 300,000 cells were incubated with 0.5 μg primary antibody or control IgG. For E14.5 and E15.5, four retinas from two animals were pooled per biological replicate; for E16.5 and later stages, two retinas from one animal were pooled per biological replicate. DNA fragments were purified on spin columns and eluted with 30 µl elution buffer (Cell Signaling Technology, Cat #14209).

Libraries were prepared by the University of Utah High-Throughput Genomics Core using the NEBNext Ultra II DNA Library Prep Kit (New England Biolabs, Cat #E7634L). PCR-amplified library quality was verified using an Agilent Technologies 4150 TapeStation with D1000 ScreenTape assay (Cat #5067-5582 and #5067-5583) and the molarity of adapter-modified molecules was defined by quantitative PCR using the Kapa Biosystems Kapa Library Quant Kit (Cat #KK4824). Libraries were normalized and pooled for paired-end sequencing (2 × 151 bp) either on NovaSeq 6000 with NovaSeq 6000 S4 reagent Kit v1.5 (Cat #20028312) or on NovaSeq X with NovaSeq X series 10B reagent Kit (Cat #20085594).

#### CUT&RUN data analysis

All CUT&RUN samples were trimmed using Cutadapt v3.5 to remove adapter sequences and then aligned to mouse genome build mm10 using Novoalign v4.02.02 or Bowtie2 v2.5.1. Samtools v1.16 was used to filter out mapped reads with less than a 30 MAPQ score, then sort, and index bam files. Unique Molecular Indexes (UMIs) were added and sequenced to identify PCR duplicates in most CUT&RUN experiments, except for H3K27me3 and H2AK119ub1 in P0 control and *Foxp1*-DisCo^-/-^ mice. The UMIs were handled by scripts written by the Huntsman Cancer Institute Bioinformatics Core, and PCR and optical duplicates were removed with optical distance threshold at 2500 (https://github.com/HuntsmanCancerInstitute/UMIScripts). For samples lacking UMIs, Picard v2.26.3 was used to mark and remove duplicates.

Downstream analysis of histone modifications was performed using MultiRepMacsChIPSeq v20.2, a MACS2 based pipeline developed by the University of Utah Bioinformatics Shared Resource (github.com/HuntsmanCancerInstitute/MultiRepMacsChIPSeq). Briefly, the pipeline used paired-end and broad peak calling mode to identify peaks for each replicate. The tolerated gap was set to 1000 bp, and the minimum q-value to 0.1. The peaks from each replicate per condition were intersected to generate consensus peaks using intersect_peaks function in the pipeline. The consensus peaks of each condition were merged, and reads under the merged peak were counted. DESeq2 v1.44.0 was used to identify significantly different peaks between conditions. All bigwig genome tracks were depth-normalized, and replicates were averaged by mean and reported as a single track. H2AK119ub1 CUT&RUN in E18.5 control and *Jarid2* condition knockout was performed in five biological replicates for each condition across two batches. The batch effect was corrected for in the DESeq2 design formula. For CUT&RUN of histone modifications across developmental stages in wild type mice, only female pups were used at E15.5 and E16.5 to match our published E14.5 and P0 H3K27me3 CUT&RUN data (5). Jarid2-PRC2.2 regulated H3K27me3-enriched regions were previously identified as having reduced H3K27me3 in *Jarid2* conditional knockout retina. Heatmaps and profile plots were generated by computeMatrix, plotHeatmap and plotProfile from Deeptools v3.5.1.

Peaks of transcription factors were called using MACS2 (v2.2.7.1) using paired-end and narrowPeak mode. Foxp1 CUT&RUN was performed in two different biological conditions. For P0 *Foxp1*-cTG samples, three biological replicates were run across two batches. Peaks were called using a minimum FDR cutoff threshold (-q) of 0.01 and peaks called in at least two replicates were identified by the intersect_peaks function in the MultiRepMacsChIPSeq pipeline. For the single E14.5 WT sample, the minimum FDR cutoff threshold (-q) was set to 0.2. The final Foxp1_consensus peaks were created by intersecting peaks identified in both P0 *Foxp1*-cTG and E14.5 WT. Peaks were annotated with nearest genes and genomic regions using ChIPseeker (v3.20) with the default settings, in which TSS (transcription start site) is defined from -3kb to +3kb. Motif enrichment was performed using findMotifsGenome from Homer v4.11.1 under default parameters.

### RNA sequencing

#### Sample and Library Preparation

RNA from P0 retinas was isolated using RNeasy Plus Mini Kit (Qiagen, Cat #74134) and cleaned up using RNA clean & Concentrator-5 kit from Zymo Research (Cat #R1013). RNA samples were processed for library generation and sequencing by the Huntsman Cancer Institute High-Throughput Genomics Core Facility. RNA concentration was measured with a Qubit RNA BR Assay Kit (Fisher Scientific, Cat #Q10211). RNA quality was evaluated with an Agilent Technologies RNA ScreenTape Assay (Cat #5067-5576 and 5067-5577). Total RNA samples (5-500 ng) were hybridized with NEBNext rRNA Depletion Kit v2 (Human, Mouse, Rat) (Cat #E7400) to substantially diminish rRNA from the samples. Stranded RNA sequencing libraries were prepared as described using the NEBNext Ultra II Directional RNA Library Prep Kit for Illumina (Cat #E7760L). Purified libraries were qualified on an Agilent Technologies 4150 TapeStation using a D1000 ScreenTape assay (Cat #5067-5582 and 5067-5583). The molarity of adapter-modified molecules was defined by quantitative PCR using the Kapa Biosystems Kapa Library Quant Kit (Cat #KK4824). Individual libraries were normalized and pooled in preparation for Illumina sequence analysis. Sequencing libraries were chemically denatured in preparation for sequencing. Following transfer of the denatured samples to an Illumina NovaSeq X instrument, a 151 x 151 cycle paired end sequence run was performed using a NovaSeq X Series 10B Reagent Kit (20085594).

#### RNA-seq data analysis

Raw reads were pre-processed with clumpify from the BBTools package (v39.06) to remove optical duplicates, with maximum distance set to 100. Reads were trimmed using Cutadapt v3.4 to remove Illumina adapters with a minimum read length of 20 and minimum overlap length between reads and adapter sequence of 6. Trimmed reads were aligned to the mouse Ensembl genome annotation release 112 with STAR v2.7.9a in two pass mode. Reads were counted with featureCounts from the subread package v2.0.3 and genes with fewer than 5 counts in every sample were eliminated. Differential gene expression analysis was performed using DESeq2 v1.32.0. Pheatmap was used to perform hierarchical clustering and plot the heatmap.

### Reanalysis of published sequencing data

E14.5 and P0 H3K27Ac ChIP-seq and ATAC-seq data were from GSE87064 (66, 67). E14.5 Atoh7 CUT&Tag data was from GSE220587 (22). Fastq files of two Atoh7 replicates were downloaded and analyzed as described in CUT&RUN data analysis. Bigwig files for IGV visualization were generated using DeepTools v3.5.6 bigwigAverage. The processed HiC data and BED files with TAD ranges from P0 mouse retina were downloaded from GSE135465 and visualized using HiCExplorer v3.7.2. Bigwig and BED files that were originally built on mm9 were converted to mm10 using CrossMap (67).

### Statistical analysis

All image data were analyzed using Prism 10 software (GraphPad, La Jolla, CA). All image quantification was tested for normality using Shapiro-Wilk test. If normality was not met, non-parametric tests were used. Homogeneity of variance was assessed using the Brown-Forsythe test. If variances were significantly different, we performed Welch’s one-way ANOVA with Dunnett’s T3 multiple comparison test. Otherwise, an ordinary one-way ANOVA with Tukey’s multiple comparison test was performed. For selected pairwise comparison, Sidak’s multiple comparisons were applied. For all data that is presented as the mean, error bars indicate the standard error of the mean (SEM). We used a 95% confidence interval, and a P value of < 0.05 was set for rejecting the null hypothesis. In all graphs, dots represent individual retinas. *p<0.05; **p<0.01; ***p<0.001; ****p<0.0001.

## Author contributions

Conceptualization: F.C. and M.L.V.; methodology: F.C., J.Z., J.M.R., J.S., J.B.M., M.C., P.M., and M.L.V.; image analysis, F.C., J.B.M., and J.S.; bioinformatical analysis, F.C. and J.M.R.; manuscript preparation, writing and editing: F.C., J.M.R., and M.L.V.; funding acquisition: J.B.M., M.C., P.M., M.L.V.

## Acknowledgement

This research was supported by the National Eye Institute award R01EY035158 (to M.L.V.) and R01EY037228 (to J.B.M.), the Canadian Institutes of Health Research FDN-159936 and PJT-195790 (to M.C.), the Canadian Institutes of Health Research PJT-166032 (to P.M), and the Natural Sciences and Engineering Research Council of Canada RGPIN-2026-05722 (to P.M). We thank the University of Utah Transgenic & Gene Targeting Mouse Core Facility for generating the CRISPR knockout mouse line. We thank the University of Utah Office of Comparative Medicine for animal husbandry. We acknowledge HSC Flow Cytometry Core Facility from University of Utah for technical support with FACS. We acknowledge support from the High-Throughput Genomics and Cancer Bioinformatics Shared Resource and HCI’s National Cancer Institute Cancer Center Support Grant P30CA042014. The computational resources used were partially funded by the NIH Shared Instrumentation grant 1S10OD021644-01A1. Sanger DNA sequencing was performed at the DNA Sequencing Core Facility, University of Utah. The content is solely the responsibility of the authors and does not necessarily represent the official views of the NIH.

